# Luminos: open-source software for bidirectional microscopy

**DOI:** 10.1101/2025.02.22.639658

**Authors:** Daniel G. Itkis, F. Phil Brooks, Hunter C. Davis, Raphael Hotter, J. David Wong-Campos, Yitong Qi, Bill Z. Jia, Madeleine Howell, Marley Xiong, Rebecca Frank Hayward, Byung Hun Lee, Yangdong Wang, Rebecca T. Perelman, Adam E. Cohen

**Affiliations:** Department of Chemistry and Chemical Biology, Harvard University, Cambridge MA 02138

## Abstract

Bidirectional microscopy (BDM) combines simultaneous targeted optical perturbation and imaging of biophysical or biochemical signals (e.g. membrane voltage, Ca^2+^, or signaling molecules). A core challenge in BDM is precise spatial and temporal alignment of stimulation, imaging, and other experimental parameters. Here we present Luminos, an open-source MATLAB library for modular and precisely synchronized control of BDM experiments. The system supports hardware-triggered synchronization across stimulation, recording, and imaging channels with microsecond accuracy. Source code and documentation for Luminos are available online at https://www.luminosmicroscopy.com and https://github.com/adamcohenlab/luminos-microscopy. This library will facilitate development of bidirectional microscopy methods across the biological sciences.

## Introduction

Combined optical stimulation and fluorescence imaging are powerful tools for studies of biological processes that are structured in space or time.^1^ Examples are: neural activity, cardiac activity, muscle activity, embryonic development, wound healing, or cancer growth. As opposed to conventional molecule-based perturbation and measurement tools, photons have the ability to be modulated in space, time, and energy (i.e. wavelength) with high resolution.^2^ One can thus use photons to probe how a perturbation delivered to one part of a sample r_1_ at a time t_1_ can affect dynamics at other parts of the sample r_2_ at later times t_2_.

There are many light-activated molecular tools which can convert a pattern of illumination into a patterned biochemical signal.^3^ For example, channelrhodopsins are light-gated ion channels which can pass an ionic current.^4^ Depending on the ion selectivity of the channelrhodopsin, that current can either activate or suppress electrical activity in an excitable cell. The blue light-activated adenylyl cyclase (bPAC) converts a blue light stimulus into production of a signaling molecule, cyclic adenosine monophosphate (cAMP).^5^ Other light-activated proteins can mimic morphogens involved in embryonic development;^6^ or can trigger growth or disassembly of cytoskeletal elements.^7^ Organic molecules can also convert patterns of light into biochemical signals. For example, azobenzene moieties can modulate the activity of a drug.^8^ Photo-caged tetrazine compounds can initiate click-chemistry reactions when illuminated.^9,10^ Photocrosslinkers can immobilize molecules or cells when illuminated.^11^

There are also many fluorescent reporters of cellular physiology.^12^ For example the GCaMP calcium indicators report Ca^2+^ ion concentration,^12^ the QuasAr^13^ and Voltron^14^ proteins report voltage, and the pink-Flamindo protein reports cAMP.^15^

To pair targeted optical stimulation and fluorescence imaging, one typically wishes to stimulate a cell with patterns of light at one wavelength, and to excite fluorescence from a reporter molecule at a different wavelength. Typically this requires combining a blue shifted optical actuator molecule with a red-shifted fluorescent reporter (since most molecules have a long blue “tail” in their excitation spectra, it is better for the actuator to be blue and the reporter red, than *vice versa*). Examples of such pairings are the “Optopatch” constructs which combine a channelrhodopsin and a red-shifted voltage indicator for all-optical electrophysiology;^13,16,17^ a construct combining bPAC and pink-Flamindo for all-optical studies of cAMP transport^18^; and a construct combining a calcium-permeable channelrhodopsin CapChR2^19^ with a far-red calcium indicator, FR-GECO1c,^20^ for all-optical studies of calcium handling.^21^

For three-dimensional samples, structured illumination imaging techniques can provide optical sectioning, reduce background fluorescence, and minimize optical heating and photodamage to the sample. Thus, BDM in tissues often requires patterning of both the light for stimulation and the light for imaging.

### Features

Luminos is designed to accommodate hardware most frequently used in bidirectional microscopy experiments (Supplementary Table 1), and to be extensible to new hardware. A detailed description of the features as of this submission is in the Supplementary Information. Up-to-date feature information is at https://www.luminosmicroscopy.com.

#### Recording

Simultaneous readout from multiple cameras, from widely used manufacturers (Hamamatsu, Teledyne Kinetix, Andor), photomultiplier tubes, electrodes, and photodiodes.

#### Hardware control

Control of digital micromirror devices (DMDs) or liquid crystal spatial light modulators (SLMs) for structured illumination; galvos for point-scanning stimulation or imaging; lasers, LEDs, acousto-optic deflectors (AOD), electro-optic deflectors (EOD), acousto-optic tunable filters (AOTF), Pockels cells and shutters, positioning stages, fluidic valves, patch clamp electrophysiology, and behavioral stimuli.

#### Registration

Fast automated registration of light-patterning devices to camera space- and time coordinate systems.

#### Virtual reality (VR)

An adapter to ViRMEn VR engine^22^.

#### Sequential experiments

Easy setup of experimental sequences with arbitrary parameter sweeps and autofocus.

#### Pre-programmed modules

Structured illumination microscopy via Hadamard^23^ or HiLo^24^ imaging; all-optical electrophysiology for mapping neural excitability or synaptic connectivity; sample-responsive stimulus protocols.

#### High-throughput acquisition

Writing to NVMe or directly to virtual RAM drive for recordings at multiple GB*/*s from multiple cameras simultaneously.

#### Graphical interface

An intuitive tab-based user interface lets non-experts perform calibrations, and load and share sophisticated measurement protocols.

#### Command-line interface

A Matlab command-line interface lets users script arbitrarily complex acquisition protocols and conditional experiment execution.

### Architecture

A detailed description of the architecture is in the Supplementary Information. In brief, Luminos consists of three software layers (Fig. 1a): The bottom layer comprises compiled C++ drivers which support performance-critical components (streaming, live video display, DAQ configuration). The middle layer is a modular MATLAB core with classes representing each device type and experiment acquisition scripts, accessible via the top layer GUI (using JavaScript with ReactJS) or the MATLAB command line. This GUI features separate tabs for different device classes along with the ability to save and load custom configurations and complex protocols. Once the software is configured for a given microscope, most experiments can be performed from this top-level interface. Upon completion of each acquisition, Luminos logs the complete state of the virtual microscope in a standardized MATLAB .mat file, guaranteeing that data from different custom microscopes can be analyzed using the same code.

**Figure 1:**
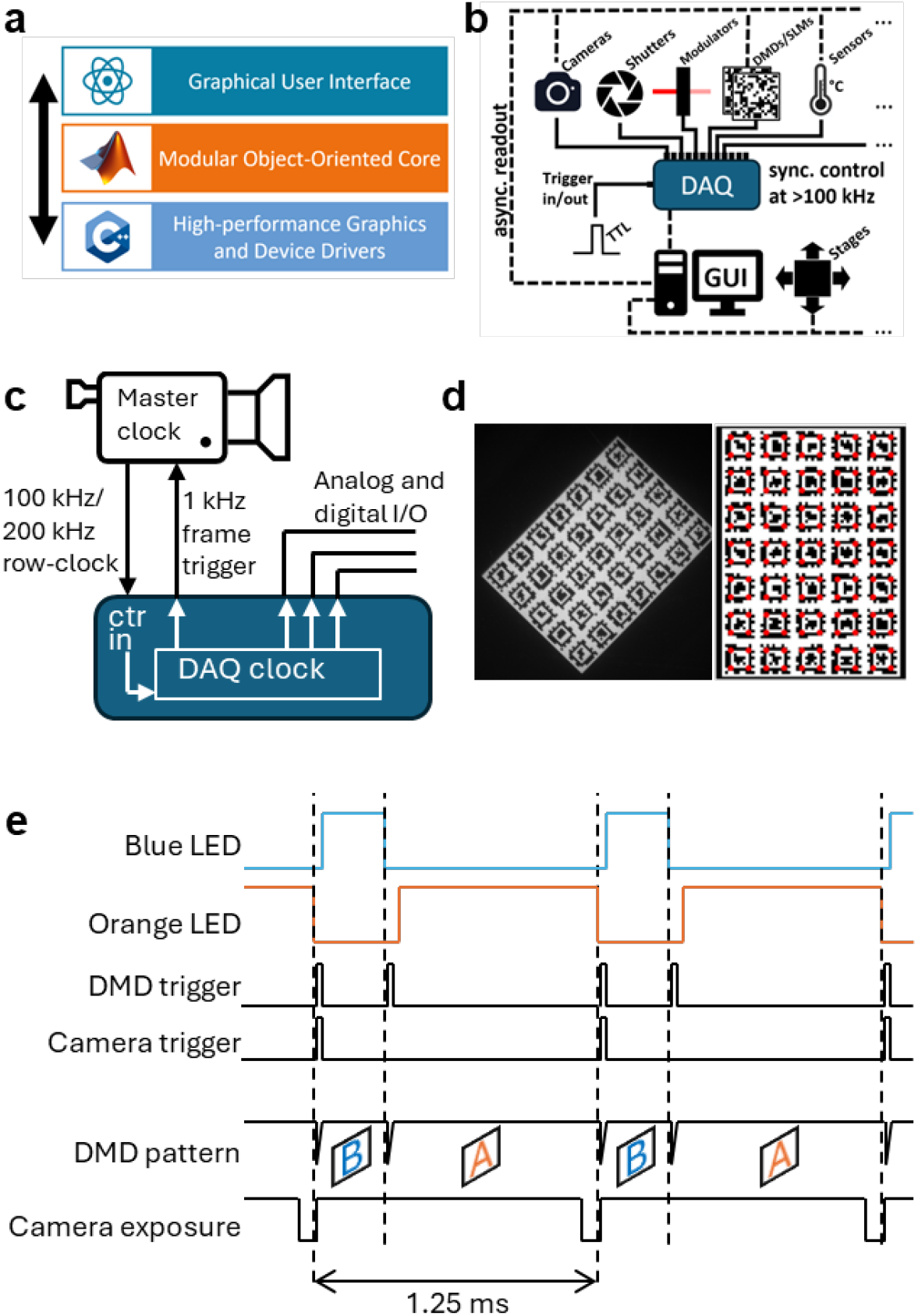
Luminos software for bidirectional microscopy. a) Luminos software architecture. b) Hardware arrangement for an experiment controlled by Luminos. c) Sub-microsecond synchronization of camera and DAQ. The camera row-clock (typically 100 kHz or 200 kHz) serves as the master clock for the DAQ. The DAQ triggers camera exposures upon integer number of row-clock pulses. All other hardware I/O functions are synchronized to the camera row-clock. d) Spatial registration of DMD and camera. A projected array of April tags is automatically read and segmented to determine mapping from DMD pixels to camera pixels. e) Sub-frame DAQ trigger scheme for multi-wavelength patterning from a single DMD.

### Hardware-software integration

Luminos is built around a National Instruments buffered data acquisition card (NI-DAQ; Fig. 1b). For an experimental run, cameras and patterning devices are pre-configured via software, and analog and digital control waveforms are loaded onto the DAQ. The acquisition is then executed synchronously, triggered by an internally or externally generated TTL pulse. The host computer sets up the acquisition, streams the data, and records the metadata, while the DAQ controls all timing, ensuring hardware-precise synchronization.

### Spatial registration

In bidirectional microscopy experiments, one typically wishes to target illumination to specific features in a sample. This targeting requires precise registration of the coordinate systems of the light-patterning and imaging devices. In Luminos, the pixels on a selected camera chip provide the reference grid to which all other coordinate systems are mapped.

The Luminos calibration function automatically finds a mapping between the patterning device coordinate system (pixel on DMD and SLM, beam offset by voltage for galvos) and the main camera. The calibration method varies by device type. For DMDs, Luminos projects an array of machine-readable April Tags^25^ onto a homogeneous fluorescent slide (Fig. 1c). Luminos automatically identifies the tag positions in a camera image and references them to the corresponding DMD coordinates. A single fully visible tag is sufficient for simple affine or projective transforms, while polynomial transforms that account for curvature around the edges of the field of view require at least 3 or 5 full tags, depending on the degree of the polynomial.

Galvos and SLMs use simpler sequential projection of individual points on a fluorescent slide or sample which are detected and registered automatically. If automatic DMD calibration using April Tags is unworkable, a manual calibration option can be selected from the calibration pattern selection, which sequentially projects individual points that are clicked manually and then refined by computational peak-finding. The specific ROI types available and any added functionality depend on the type of patterning device.

### Timing registration

In Luminos, the DAQ is the master controller of all hardware-triggered elements. National Instruments DAQ cards have high-precision internal clocks, which can provide sub-microsecond synchronization of multiple analog input/output (I/O), digital I/O, and counter operations. For Hamamatsu scientific CMOS (sCMOS) cameras operated in “synchronous” triggering mode, the camera sets the exposure time by the interval between upstrokes of a TTL frame-trigger signal. In this case, a potential problem arises at high frame rates. Scientific CMOS cameras have an internal row-clock which synchronizes the readout. These clocks are typically 100 kHz or 200 kHz. When an exposure trigger arrives part-way through a row-clock cycle, the camera waits until the end of the row-clock cycle (up to 5 or 10 μs) to begin the exposure. The asynchrony of the DAQ and camera clocks introduces a jitter in the exposure time of 5 or 10 μs. While normally unimportant, on a 1 ms exposure time, this jitter corresponds to 1% noise, a substantial noise source for voltage-imaging experiments.

In response to this issue, Hamamatsu provided user access to the internal row-clock. The synchronization problem is addressed by using the camera row-clock as the master clock for the DAQ (Fig. 1d). With this approach, the DAQ outputs retain a fixed phase relationship to the camera clock cycles, and the jitter is eliminated. In the Teledyne Kinetix camera, the frame duration is pre-specified in software. Provided that the interval between frame triggers from the DAQ exceeds the frame duration, all frames have identical exposure times and it is not necessary to use the camera as the master-clock for the acquisition.

### Multi-wavelength patterning from a single DMD

BDM experiments often involve stimulating a sample with one pattern of illumination, and then mapping a response via either wide-field or structured illumination microscopy. In these cases, one typically wishes to have distinct patterns of illumination for two (or more) wavelengths. This goal can be achieved with a digital micromirror device (DMD). To display two concurrent patterns, one can use two separate DMDs and then combine the images from each, or split a single DMD into two sub-regions and pattern a different wavelength on each^16^. However, this approach is costly, is limited to two concurrent patterns or wavelengths, and requires careful registration and co-focusing between patterned wavelengths.

An alternative is to interleave two or more patterns on the same DMD while synchronously modulating the illumination wavelengths (this is how color images are created on most commercial video projectors). Provided that the response kinetics of the light-sensitive molecular actuators are slower than the cycling frequency of the DMD, the light-responsive molecules effectively experience the time-average illumination. To implement this approach, the images displayed on the DMD must be precisely registered in time and space with the images acquired on the camera.

Fig. 1e shows how Luminos uses a single DMD to pattern multiple wavelengths. Commercially available DMD modules can switch between pre-loaded patterns in < 100 μs, permitting time-sharing of a single camera frame between two or more illumination patterns. Synchronous switching between DMD patterns during each camera frame avoids aliasing of light modulation noise into the camera measurement band. Synchronous modulation of illumination wavelength and DMD patterns completes the protocol.

### Comparison to other software

Microscopy control software packages vary in their implementations and features^26^ (Fig. 2). Hardware-timed synchronization in Luminos differs from μManager^27^, which uses the computer operating system for timing and thus encounters variable delays, up to tens of milliseconds. Luminos has advanced tools for camera-based 1P microscopies, while ScanImage^28^ or Scope^*^ are optimized for point-scanning microscopies.

**Figure 2.**
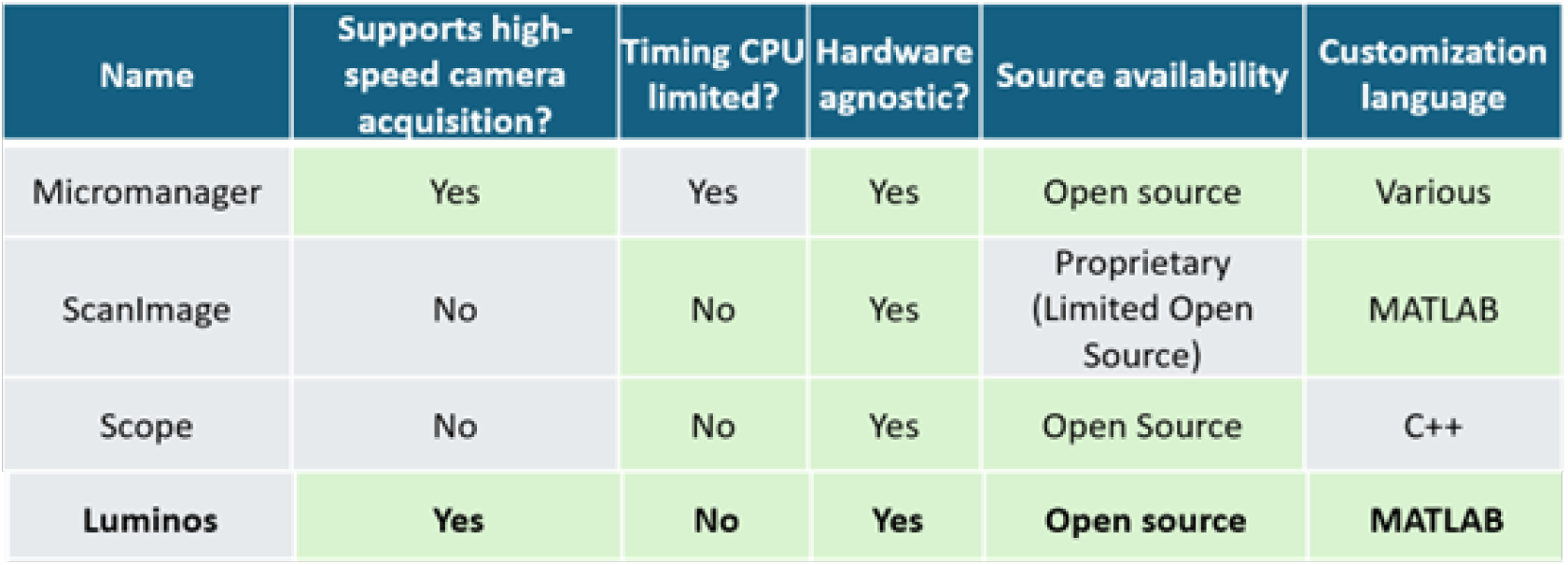
Comparison of Luminos to other microscope-control software. Luminos provides unique features for precisely registered patterned illumination and high-speed imaging.

### Applications

Representative uses include: Targeted optogenetic perturbations and high-speed structured illumination voltage imaging to map bioelectrical responses of neuronal dendrites (Fig. 3a) ^29,30^, functional connectivity mapping in neural circuits *in-vivo*^31^, optical control of morphogen signaling in developing embryos^32^ (Fig. 3b), and optogenetic stimulation and calcium imaging to map zebrafish heartbeats^33^ (Fig. 3c).

**Figure 3.**
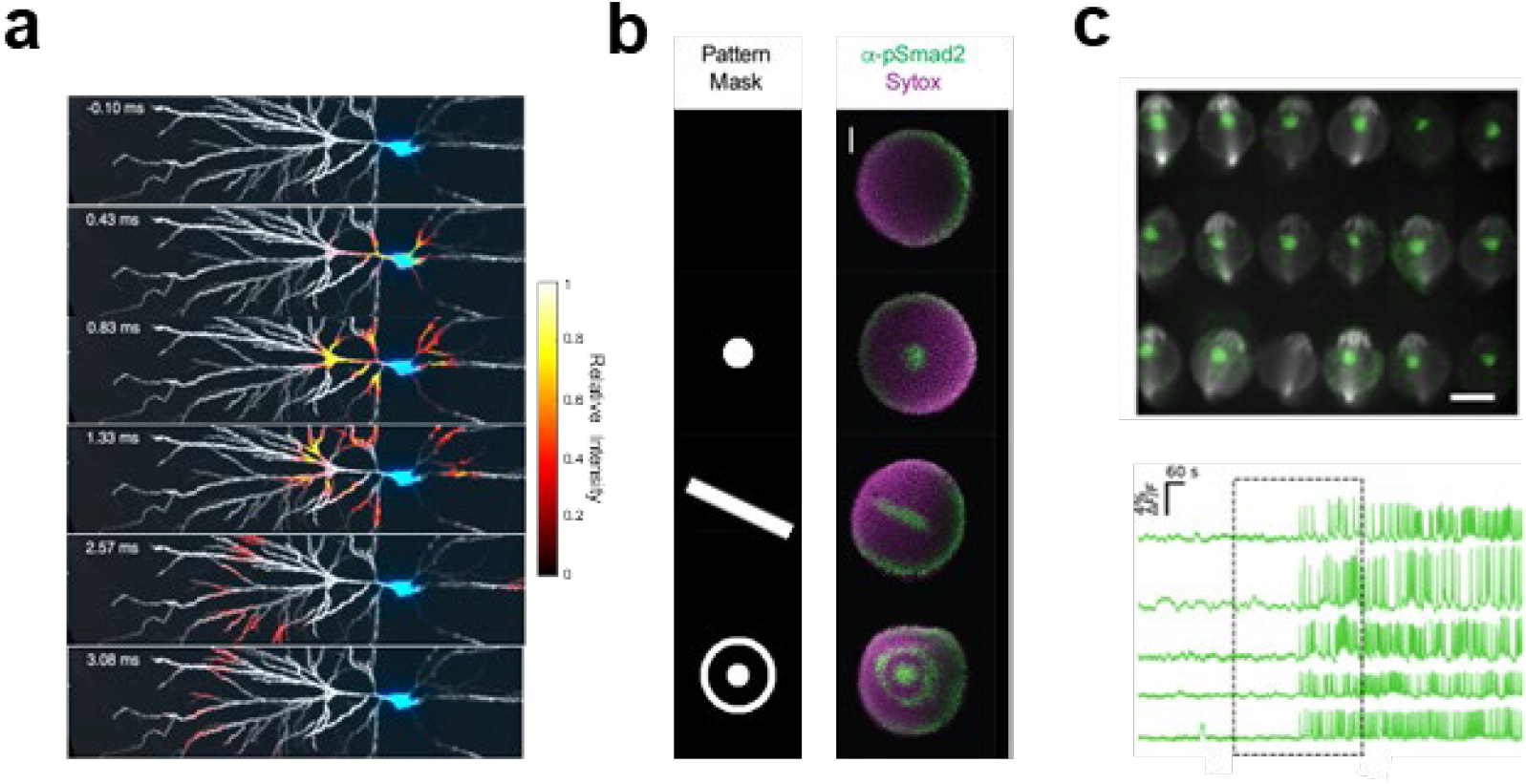
Example applications of Luminos. a) Soma-targeted optogenetic stimulation of a CA1 pyramidal neuron evoked a backpropagating action potential whose wavefront was mapped via Voltron2^29^. b) Optogenetic perturbation of Nodal signaling in Zebrafish embryos using optoNodal2, with α-pSmad2 immunostaining (green) demonstrating spatial patterning of signaling activity^32^. Scale bar 100 μm. c) Top: Zebrafish embryos with green jGCaMP7f calcium indicator in developing heart. Scale bar 500 μm. Bottom: fluorescence from five embryos, aligned by the first heartbeat^33^.

## Acknowledgments

We thank all members of the Cohen Lab, Alec Barrios and Yangluorong Liu for feedback on Luminos. This work was supported by Chan Zuckerberg Initiative Dynamic Imaging grant 2023-321177, a Vannevar Bush Faculty Fellowship, and NIH grants R01-NS126043, R01-NS133755, and R01-MH117042.

## Footnotes

* http://rkscope.sourceforge.net/.

## Notes

### Competing Interest Statement

The authors have declared no competing interest.

