## Supplemental Information for "Luminos: open-source software for bidirectional microscopy"

### Supplementary Information

#### Luminos architecture

Luminos' core is implemented in modular object-oriented code in Matlab and C++.  A ReactJS layer provides a tab-based graphical interface.

The top-level unit of function of Luminos is an instance of a Rig_Control_App object. The Rig_Control_App superclass manages device initialization, JavaScript communication, experiment coordination, and saving of metadata. Distinct subclasses of the Rig_Control_App are made for each microscope, and multiple subclasses (virtual microscopes) can share overlapping sets of hardware. Microscope-specific subclasses may implement additional functionality.

Individual hardware devices or virtual devices (e.g. a single analog output from the DAQ) are represented by instances of the abstract Device superclass. We implement a multilevel class hierarchy so that, for example, functionality common to all types of patterning devices (calibration, ROI selection, etc.) is specified in a mid-level abstract Patterning_Device superclass, with the particular implementations given in device-specific subclasses. This allows for device-agnostic modular code so that, for example, a DMD patterning tab can be implemented without knowing which specific DMD device will be used.

Each device class consists of: properties that are automatically included in the metadata archived after an acquisition; transient properties that are not saved; and methods that implement the device functions. Device class instances are independent of the Rig_Control_App that loads them and of other devices and may be loaded without loading the app using the Standalone_Device() utility. This ensures that devices may be loaded in any order and maintains control over inter-device communications at the app level for more transparent and robust code.

Luminos uses modular C++ code for performance-critical device implementation that benefits from thread parallelization or lower-level device access. We currently implement camera and live video streaming, DAQ communications, galvo scanners, and ALP DMDs at this level. We use hierarchical class inheritance to specify a common interface in a high-level class while providing the device-specific implementation in a subclass. The high-level Cam_Wrapper class is currently subclassed by Hamamatsu_Cam, Andor_Cam, and Kinetix_Cam. We split the compiled code into two functional parts. The custom device drivers are compiled to C++ object files, which allows testing via compiled C++ testbenches. An adapter layer implements the MATLAB interface and is compiled into mex files which provide a library interface to MATLAB. This separation of the interface from the driver implementation allows use of these drivers with a different high-level language, e.g. Python, simply by writing a different adapter layer. We implement optional optimized vectorized instructions for parallel computation using SSE 4.1 vector intrinsics on CPUs that support this (Intel after Penryn 2007/8; AMD after Bulldozer 2011). To maintain cross-hardware compatibility, we do not implement any GPU-based computations.

The ReactJS user interface comprises a series of tabs. Each tab communicates with a set of devices, implements user device control and inter-device coordination, and may contain subpanels for specific sub-functionality. A tab may also communicate with components of other tabs. All apps will have a main tab (Suppl. Fig. 1), which contains several subpanels, and a waveform tab (Suppl. Fig. 2). Other tabs are optional, and custom tabs can be created using the ReactJS framework. The user interface communicates with Matlab via a local Node.JS server.

**Luminos information flow**

There are two flows of information controlled by Luminos, asynchronous device configuration and synchronous device control (Fig. 1b).

The flow of asynchronous device configuration information begins with the rig initializer files. Each microscope app has a corresponding rig initializer file in JSON format. Upon initialization, the Rig_Control_App instance loads this initializer and initializes each device represented in the file with the corresponding properties given in the file. This initializes the microscope into a consistent predefined state. After initialization, asynchronous commands can be sent either from the user interface or from MATLAB to update device configurations. At the end of an acquisition, all devices that have been flagged for monitoring (using the Rig_Control_App.assignDevicesForMonitoring() method) will have all non-transient properties saved into the output_data.mat metadata file for use in analysis. Patterning device calibrations are also separately saved in MATLAB .mat files, which are updated upon calibration. This allows calibration to be maintained between imaging sessions. Asynchronous device communication is implemented in a device-specific manner. Many devices implement ASCII serial commands, while others come with driver libraries.

The flow of information for synchronous device control begins on the waveform tab, where custom waveforms are defined. These waveforms can be saved and loaded for reuse. Before acquisition, the waveform is built and loaded into the DAQ buffer. Upon triggering, the DAQ begins synchronous buffered operation of all defined outputs and inputs until the predefined acquisition time is complete. Upon completion, the input and output buffers are saved with corresponding metadata into the output_data.mat file. DAQ operation can be clocked either internally or externally. All synchronous control is implemented as analog and digital input-output links between the DAQ and the hardware devices.

For precisely timed camera acquisition, it is often necessary that the camera and DAQ share the same clock. Many modern cameras, including Hamamatsu Flash and Fusion and Andor iXon cameras, provide a clock output that can be used to clock the DAQ. Similarly, for two-photon scanning systems, it is possible to clock the DAQ from the pulsed laser. It is not possible to ensure a shared clock if there are multiple devices that each run only on an internal clock.

*Configuring and using RAM-drive mode in Luminos*

Modern NVMe SSD drives and/or RAID SSD arrays can support sufficient data throughput to record camera data at high speed directly onto disk. Systems with slower storage must stream into RAM instead.

Assuming sufficient RAM is installed to store a full acquisition while still leaving sufficient RAM for program execution, Luminos can be configured to stream high-speed video directly to RAM, copying the result over to the Data disk directory after the acquisition finishes. To do this, first install ImDisk Toolkit (https://sourceforge.net/projects/imdisk-toolkit/) and create a RAMdisk of the necessary capacity with drive letter R:. Then set the "rdrivemode" parameter to 1 in the camera device entry of the app initializer (see tutorial on initializers below). Acquisition will now stream to RAM and will not be limited by disk storage speed. We do not enable RAM storage by default because the added transfer step can be unreliable. We recommend fast disk storage when possible.

**User interface**

A detailed description of every function in the user interface is available at:
https://www.luminosmicroscopy.com

**1. *Main tab:***
The Main tab (Suppl. Fig. 1) is required for camera acquisition and contains a series of camera-related and general utilities:

Experiment name: Text entry field for naming each acquisition.

Toggles & modulators: Panel that provides real-time control over all digital shutters, analog modulators, and filter wheels.

Stage controller and z-stage controller: For systems with a computer-controlled stage, a stage controller panel provides control and readout of current stage position.

Notes: User notes saved into the user directory in "session_notes.txt".

**1.1 Camera Controls:** When multiple cameras are present, the camera control panel will also be tabbed, with a tab for each camera.

Start Acquisition: Starts camera recording with parameters set in Main tab and the waveforms set in the Waveform tab. This calls the script Waveform_Camera_Sync_Acquisition.m, which can be modified according to individual needs. During recordings, Luminos sets a series of timers to wait for the end of the acquisition, without blocking access to the MATLAB terminal or the execution of other scripts. Once the DAQ has finished running its waveforms, acquisition is finished, cameras are restarted (whether there are outstanding requested frames or not) and "Done with experiment." is printed to the MATLAB terminal.
Cancel Acquisition: Abort active acquisition at any stage and resets running DAQ waveforms and cameras.
Blank Screen: An option for blacking out all screens during data acquisition avoids optical noise from monitor glare. This mode can be exited by pressing any key; and the monitors un-blank automatically at the end of the recording.

Exposure (s): Set camera exposure time. Minimum exposure time in seconds should be selected based on size of selected ROI and camera's line readout rate. Selecting a shorter exposure time than the hardware and ROI-specific maximum will lead to dropped frames.

Frames: Number of requested Frames can be entered manually or using the Auto button.

Auto button: The Auto button calculates maximum number of frames for the selected Duration in the Waveforms tab (Duration/Exposure time). When using Manual triggering in Frame Triggering Selection, Auto will automatically calculate the number of rising edges to be sent to camera trigger based on the waveforms defined in the Waveforms tab.

Snap: The Snap button takes a snap and saves it to the user's directory in the Snaps folder as a .tiff file, alongside a .mat file containing metadata about the snap (binning, ROI, DMD tform during snap, etc.).

Rotate FOV and flip FOV: Rotate FOV buttons will rotate the camera stream by 90 degrees each time they are clicked, while Flip FOV flips stream horizontally. These transformations are only a convenience for viewing and have no effect on the saved image data or snaps.

ROI settings: Select between arbitrary ROI (define ROI as top-left corner and height/width), centered for ROI in the center of the camera (as some camera types always start reading out from the sensor midline), or centered with offset (to adjust width and height of FOV without modifying top-left corner).

Binning: Binning sums the images 1 pixel, 2x2 pixel or 4x4 pixel on the hardware level. Selecting binning does not require any changes to ROI as ROI is always defined with respect to physical sensor pixels.

Frame triggering: *See Luminos manual for a detailed discussion of triggering modes*. The frame triggering mode dropdown allows configuration of the camera to synchronize each frame to a trigger pulse provided by the DAQ. Through the use of different trigger settings and DAQ ports, Luminos supports a range of triggering options: The "Single Start Trigger" mode sends a single pulse to the camera at the start of the recording, after which the timing and duration of the remaining frames are controlled by the camera's internal clock. "Trigger each Frame" requires a DAQ trigger to initiate or finish each frame, depending on the camera model. The DAQ trigger frequency can be set in an input field that is conditionally displayed when this mode is selected. To run the camera at a constant frame-rate in this mode, the DAQ uses a counter source/output port pair to provide regular pulses synchronized to the camera row-clock. To achieve a variable frame-rate (e.g. bursts of high-speed acquisition separated by rest periods), Luminos can provide an arbitrary binary waveform output using the "Manual Setup in Waveforms" option.

Camera ROI means plots: Displays one real-time plot per camera displaying average intensity in a user-selected region of interest.

Objectives & tube lenses: A utility for calculating magnification depending on the objective and tube lens allows for specification of pattern sizes and measurement of distances in absolute units.

Exit Button: The app exit button is present on all tabs and allows clean exit from the app. Upon exit, the app shuts any open shutters, sets analog outputs to zero, and, if configured, automatically copies all data from that session to a pre-specified remote server directory that is independently configured for each user.

**2. *Waveform tab*:**

An experimental sequence is made up of one or more analog outputs, digital outputs, and analog inputs, each of which is called a "waveform". The waveform tab (Suppl. Fig. 2) configures buffered waveforms that will be loaded onto and run from the DAQ as a single experimental sequence.

Plots: The plot at the top provides a preview of the waveforms.

Global properties: The global properties panel sets overall waveform duration, DAQ clock rate and source, the trigger mode and the optional output trigger. Options for the clock source include the internal DAQ clock or a clock connected to any of the DAQ's PFI ports. The clock rate can be set arbitrarily when using the internal DAQ clock or non-camera clock source (in which case it must match the external clock rate), while camera row-clock rates are read from the JSON initializer file and set automatically when using a camera as clock source.

Analog/digital outputs and analog inputs: Pre-defined waveform function files provide an expansible library of customizable waveform types. Individual inputs or outputs are added using the respective plus buttons, and removed using the respective minus buttons. Complex analog or digital output waveforms can be built by assigning multiple single-valued waveforms to the same output port. All waveforms sharing an output are multiplied together, and the resulting product waveform is displayed in the preview at the top of the tab. Waveform configurations can be saved and loaded with buttons at the top right. A waveform-only acquisition with metadata collection but without video can be started with another button, and can be aborted from the main tab.

**2.1 DAQ connections**

Luminos can use several optional connections to the DAQ to control waveform execution. These are:

Clock connector (digital PFI I/O): Accepts row-clock from a camera (typically 100 kHz or 200 kHz) to serve as master-clock for the DAQ, as an alternative to the DAQ's internal clock.

Start Trigger connector (digital PFI I/O): Accepts or provides a digital pulse (depending on the Trigger Mode) to signal the start of an acquisition. The start of an acquisition can be indicated in two ways:

- Self-trigger mode: DAQ starts upon software command and sends a trigger pulse out on the trigger connector.
- External-trigger mode: DAQ begins acquisition when it receives a pulse on the trigger connector.

In Self-Trigger mode, the DAQ sends a pulse to this port to initiate waveforms. The connector can remain unconnected in this case. In External Trigger mode, Luminos will prepare all devices for the acquisition and await a trigger input to this connector from an external source.

Output Completion trigger (digital non-PFI I/O): The output trigger is sent after the completion of an acquisition run, e.g. to trigger an autosampler to move to the next sample.

**3. *Camera viewer tab*:**

Every camera has a separate video live stream associated with its underlying C++ instance (Suppl. Fig. 3). This window displays the camera field of view (FOV) in real time along with a scale bar with absolute pixel values and a histogram. Using keyboard shortcuts (summarized in the main tab), the stream window can be used to execute functions such as: select ROIs, zoom in/out, change the color map and range (switch between automatic or manually set fixed, adjust fixed scale) or set binning. Optional histogram equalization provides a nonlinear adjustment to the colormap to assign approximately equal numbers of pixels to equally spaced contrast bins. This transformation maximizes visual contrast. Distances in camera pixel (or absolute length if the magnification is set) can be measured by right-clicking on any two points in the FOV.

**4. *Light patterning tabs*:**

All light patterning device tabs (Suppl. Fig. 4) allow loading and display of a reference image, a calibration button, and ROI selection tools. The DMD tab allows creation of polygonal, circular, and freeform ROIs.

Multiple ROIs can be combined (Suppl. Fig. 4). A patterning device calibration tab provides options for automated or manual mapping of patterning-device coordinates (e.g. DMD pixels) to camera pixels (Suppl. Fig. 5).

**5. *Multi-round tab*:**

The Multi-round tab (Suppl. Fig. 6) is used to set up and run multi-round experiments after setting waveform and camera parameters in their respective tabs. This tab also has options for specialized imaging techniques (Hadamard or HiLo imaging). Start and completion triggers are expected and sent at the start and end of every round of the multi-round experiment. All multi-round experiments are also directly callable from the MATLAB command line.

**5.1 Experiment selection:**

<u>Number of experiments:</u> Sets number of repetitions for selected experiment.

<u>Experiment types:</u> This drop-down menu offers options for different experiment types. Currently, option include standard acquisition, waveform only acquisition, snap only, Hadamard, and HiLo. Hadamard and Hilo are structured illumination microscopy techniques which require a DMD and only appear if a DMD device is found in Luminos. If multiple cameras or DMD devices are available, additional menus for device specification appear for snap only, Hadamard and HiLo imaging. For Hadamard and HiLo, DMD patterns and DAQ waveforms are generated and executed automatically according to the camera exposure time.

<u>Run experiments in loop:</u> Runs selected experiment type for the specified number of times.

**5.2 Scan parameters:**

The multi-round tab allows the definition of an arbitrary number of scanned parameters, that is, parameters that are updated between subsequent runs. New scan parameters can be added by clicking the plus button and removed using the individual minus buttons and are executed from top to bottom based on the order in the UI before the start of each experimental round.

There are three available types of scans:

- Linear scans, which linearly interpolate values between a start and end value using linspace.
- Custom scans, which cycle through a set of explicitly defined comma-separated values.
- Autofocus, only available for xyz-Stages and z-Stages, refocuses between acquisitions by finding the position within selected Search Range (in µm for xyz-Stages, mm for z-Stages) which maximizes 99.99th percentile pixel brightness or the highest 5% of spatial frequencies in the image. This scan type takes a "search range" and "Refocus once every" as additional inputs. 100 µm of Search Range will search from 100 µm below to 100 µm above current position, using 2 quick scans (one coarse, one fine around maximum of coarse scan). This usually takes ~6 s, depending on the search range and exposure time. This is particularly useful for cell culture and other thin samples. "Refocus once every" sets how often to refocus, e.g. 3 will refocus once every 3 acquisitions. This is compatible with all experiment types.

  Autofocus requires some light to be on, but modulators and shutters are turned off or closed between acquisitions. To use autofocus, a simple waveform with constant values for each shutter and light source and modulator that needs to be open (duration and trigger settings do not matter) can be defined in the Waveforms tab, and saved as "Autofocus.json". The Autofocus function will select the most recent file containing “Autofocus” in its title and will open shutters and set modulators as in this file for the duration of the autofocus process.

Parameters that can be scanned between rounds of experiments are:

- Motorized Z/XY/XYZ stage positions: Stages with a z-component support three types of scanning, XY-stages only “Linear Scan” and “Custom Scan”. An additional “Use current” button loads and inputs current stage position for start or end values or automatically adds the current stage position to list in the right format to “Custom Scan” positions. This scan type is useful for z-scans or for sequentially imaging multiple FOVs in a large sample or a multi-well plate.
- DMD patterns: If a DMD device is connected, a selection of patterns from a pattern stack in dmd.all_patterns can be scanned through. The only scan type here is “Custom Scan”. In the example in Suppl. Fig. 6, the first experiment sends patterns 1, 2 and 3 to DMD, then 1 and 2 for the second experiment, and then the software alternates between these. Switching patterns requires DAQ trigger pulses to best sent to the DMD trigger port. A preview window and slider are available for each DMD, to examine the stack in dmd.all_patterns and to select which patterns to add. Patterns are automatically added when using “Export” function (Chevron pattern button) in the DMD tab or the “Generate Patterns” in the Hadamard setup dialog, or can be added manually. This scan type is useful for stereotyped experiments requiring a sequence of DMD patterns, such as round-robin-style functional connectivity mapping.
- Analog and Digital waveform parameters: After the waveforms have been set in Waveforms tab, all waveform parameters appear as scannable parameters. The only scan type is “Custom Scan”. Using this overwrites the value set in the waveform tab and cycles through the custom values instead. This scan type can be used to systematically vary e.g. stimulation intensity or frequency.

**Command-line interface**

While the ReactJS user interface is the most intuitive way to configure an experiment, all functionality is present in the MATLAB layer. Devices and experiments can therefore be directly configured from MATLAB using scripts or the command line (Suppl. Fig. 7). This option allows arbitrarily complex experiments, e.g. comprising nested loops or conditional execution of protocols. Configuration of cameras and DAQ waveforms and initiation of all acquisitions is performed by experimental scripts that are triggered by the user interface acquisition buttons. These scripts set up the data directory, set up any necessary advanced timing, configure the master device that will define the end of an experiment, and send software triggers to the DAQ and camera.

**Tutorials**

The following tutorials cover initial setup of Luminos and basic customization. For additional information, consult the Luminos website https://www.luminosmicroscopy.com/, which hosts additional tutorials and a user manual, and GitHub https://github.com/adamcohenlab/luminos-microscopy.

*Tutorial 1: Initial setup*

1. Install prerequisites
    a. Windows 10/11
    b. MATLAB r2021a or later (tested up to r2024a) with the following toolboxes
        i. Data Acquisition Toolbox
        ii. Image Processing Toolbox
        iii. Instrument Control Toolbox
        iv. Optimization Toolbox (Optional)
        v. Statistics and Machine Learning Toolbox (Optional)
    c. Install the Data Acquisition Toolbox Support Package for National Instruments NI-DAQmx Devices from the MATLAB Add-Ons Manager.
    d. Node.js (https://nodejs.org/en/download/). Default Options
    e. Visual Studio Community (https://visualstudio.microsoft.com/). Desktop C++ development.
2. Clone the Luminos repository by running this line in your terminal
    a. git clone https://github.com/adamcohenlab/luminos-microscopy.git
3. Build the libraries by running the following command in MATLAB from the Luminos directory
    a. build
4. Plug in your NI DAQ or set up a DAQ simulator (https://knowledge.ni.com/KnowledgeArticleDetails?id=kA03q000000x0PxCAI&l=en-US) and note the name (You can use NIMAX to view this easily). e.g. “Dev1”
5. Edit the initializer file
    a. Open Simulator.json (type “edit Simulator.json” in MATLAB) and replace all instances of “Dev1” with the name of your DAQ device.
6. Run the app in MATLAB with the following command
    a. app = Simulator_App()
7. You should now see a simulated camera with diagonal stripes along with a browser-based simulator UI. Follow the next tutorial to configure Luminos for your hardware.

*Tutorial 2: Customizing an initializer for your hardware*

1. You will be modifying the Example_App.m and Example.json files for this. You can later rename the files and appropriate names in the files to any app name you like.
2. Open src\Applications\Example\Example.json in a text editor
3. First, edit the “dataDirectory” field to a directory where you’d like Luminos to save your data. Luminos will create timestamped subfolders here for each experimental session.
4. Now edit the “tabs” list with one entry for each tab you want. You’ll need at least “Main” and “Waveforms”. Other options are “DMD”, “SLM”, “Scanning”, “Lasers”, “Hadamard”, “SpinningDisk”. (\src\User_Interface\frontend\src\tabs for a full list). Order does not matter.
5. Now, set up the “DAQ” and “Camera” device entries with the appropriate parameters (see notes on specific options in next section).
6. Add or modify any other device entries you need, following the examples given in Example.json. The required parameters for each device can be found in the corresponding initializer. For instance, if you want to know the parameters you should specify for a given deviceType, inspect the file called <deviceType>_Initializer.m and look at the properties the initializer expects. A list of supported devices is given in Supplementary Table 1.

7. Add required device drivers.
    a. Since we are not permitted to distribute proprietary device drivers with our code, you will have to install any required drivers yourself (e.g. for cameras).
    b. For Hamamatsu cameras, download both DCAM-SDK and DCIMG-SDK (https://dcam-api.com/sdk-download/). You'll need to create a free account. Unzip both SDKs into /src/lib/Luminos_VS/inc/. You should have the following folders:
        i. src/lib/Luminos_VS/inc/dcamsdk4
        ii. src/lib/Luminos_VS/inc/dcimgsdk
    c. For Andor cameras, download the appropriate Andor SDK (https://andor.oxinst.com/downloads/) and unzip into /src/lib/Luminos_VS/inc/.
    d. After adding all drivers, recompile the C++ code by running the "build" command in MATLAB.
8. Test each device
    a. After adding each device, test by running in standalone device mode. For example, the camera would be tested with:
        i. cam = Standalone_Device("Example","Camera");
9. Run the full app
    a. After adding and testing devices separately, run the full app with:
        i. app = Example_App();
10. View the data
    a. After an acquisition, you should see in your data directory a date-stamped subfolder with a time-stamped subfolder for each acquisition.
    b. Within an acquisition folder you should see two files. The video is stored in frames1.bin, and the metadata in output_data.mat.
    c. From MATLAB, you can load the results by running the following command:
        i. [mov, avgImg, DeviceData] = Extract_Mov("path/to/data_directory");

**Supplementary Table 1: Currently supported hardware**

| | |
|---|---|
| **Software Requirements** | |
| | Windows 10/11 with MATLAB r2021a or later |
| **Camera** | |
| | Hamamatsu (tested on Flash, Fusion) |
| | Andor (tested on iXon) |
| | Teledyne Kinetix |
| **DAQ** | |
| | National Instruments (tested on USB, PCIe) |
| **DMD** | |
| | ViALUX ALP 4.1, 4.2, 4.3 |
| | TI DLP |
| **SLM** | |
| | Meadowlark |
| **Scanner** | |
| | Any analog galvo scanner |
| **Filter Wheel** | |
| | Thorlabs |
| | Optec high speed filter wheel |
| | ASI FW1000 |
| **Shutter** | |
| | Any analog or digital-controlled shutter |
| **Modulator** | |
| | Gooch & Housego 8-channel AOTF controller |
| | Any analog-controlled modulator |
| | Thorlabs motorized rotation mount with half-wave plate |
| | LED drivers |
| **Motion Control** | |
| | Sutter MPC200 |
| | Scientifica SliceScope |
| | Ludl MAC5000/6000 |
| | Thorlabs TCube (stepper and servo) |
| | Thorlabs MCM301 |
| | Newport Linear Motion controller (TRB25CC) |
| **Power Meter** | |
| | Thorlabs PM400 |
| | Newport 84x_PE |
| **Laser (software control optional)** | |
| | OBIS |
| | Hubner |
| | SpectraPhysics DeepSee |
| **Spinning Disk** | |
| | Yokogawa |
| **Optical Trap** | |
| | Any analog galvo-based trap |
| **Optical Parametric Amplifier** | |
| | Amplitude Systems Mango |
| **Other** | |
| | Any device compatible with DAQ I/O |

### Supplementary Figures

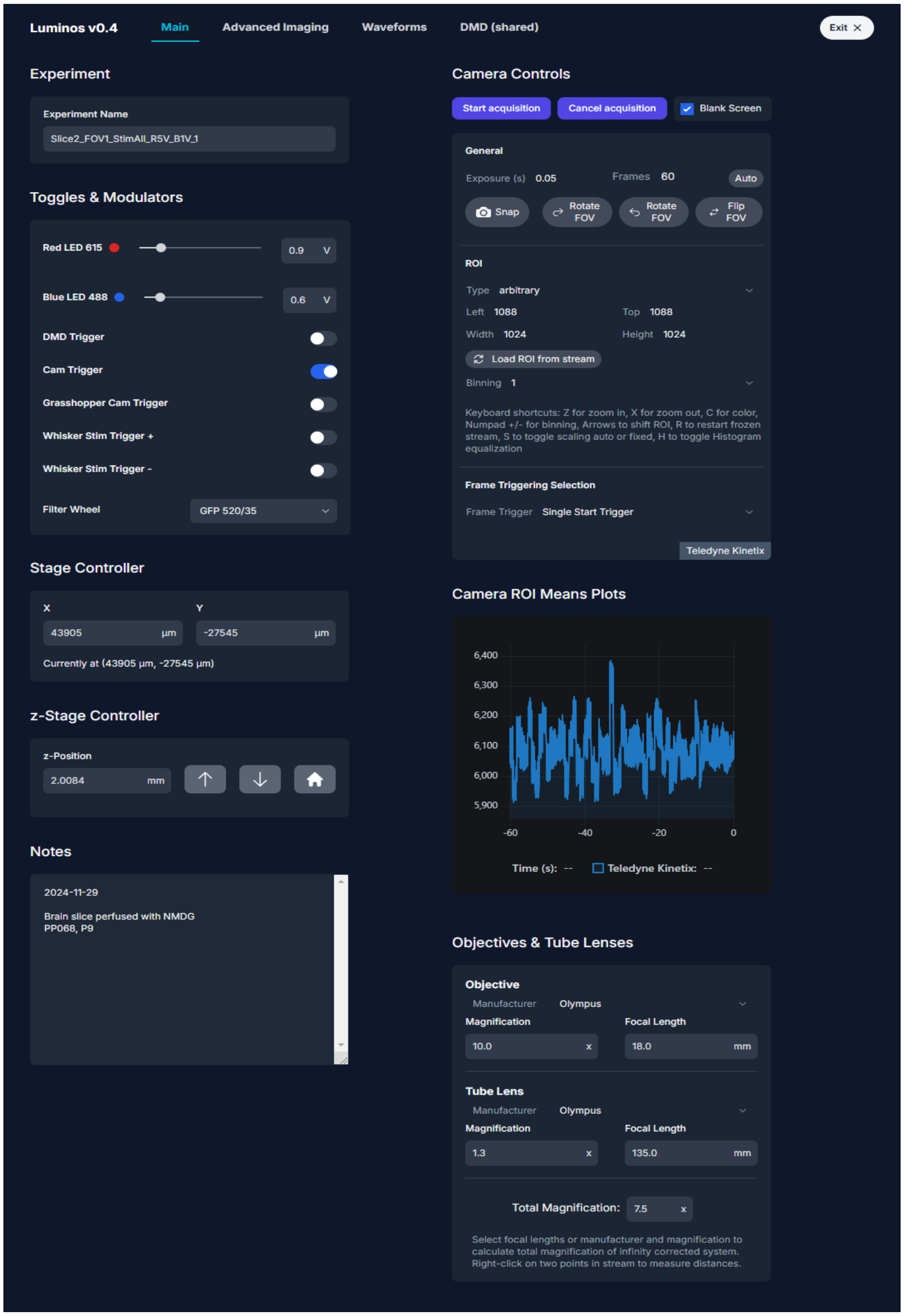


**Supplementary Figure 1: Luminos main tab.** The main tab provides overall experiment control, with experiment naming, on-demand control of analog modulators, digital shutters, filter wheels, motion control, and control of the camera. A live-updated plot shows a trace of the brightness in a user-selected region of interest from the live camera stream. A notes panel automatically saves any user notes. The tab bar at the top provides access to the other tabs. Objectives panel allows for quick calculation of magnification and calibrates distance measurements in the camera stream.

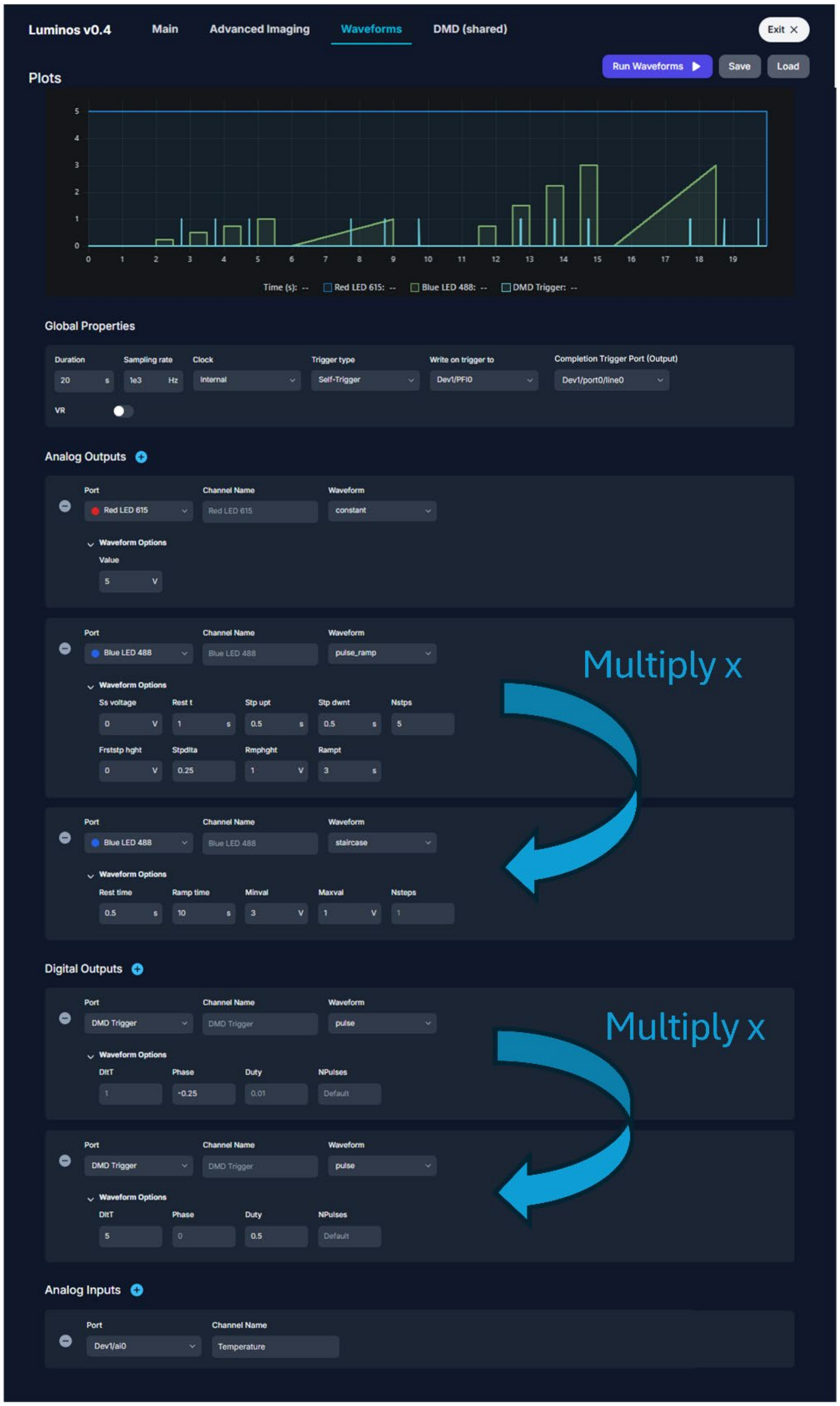


**Supplementary Figure 2: Luminos waveform configuration tab.** The waveform tab species custom buffered waveforms with the ability to save and load configurations. Clocking and triggering options provide control over timing, and the plot gives a preview of the waveform. The dropdown interface provides control without clutter. Complex waveforms are set up by multiplying primitives, or waveforms can be defined separately and loaded. When an external clock is used, the sampling rate must be set to match the external clock rate.

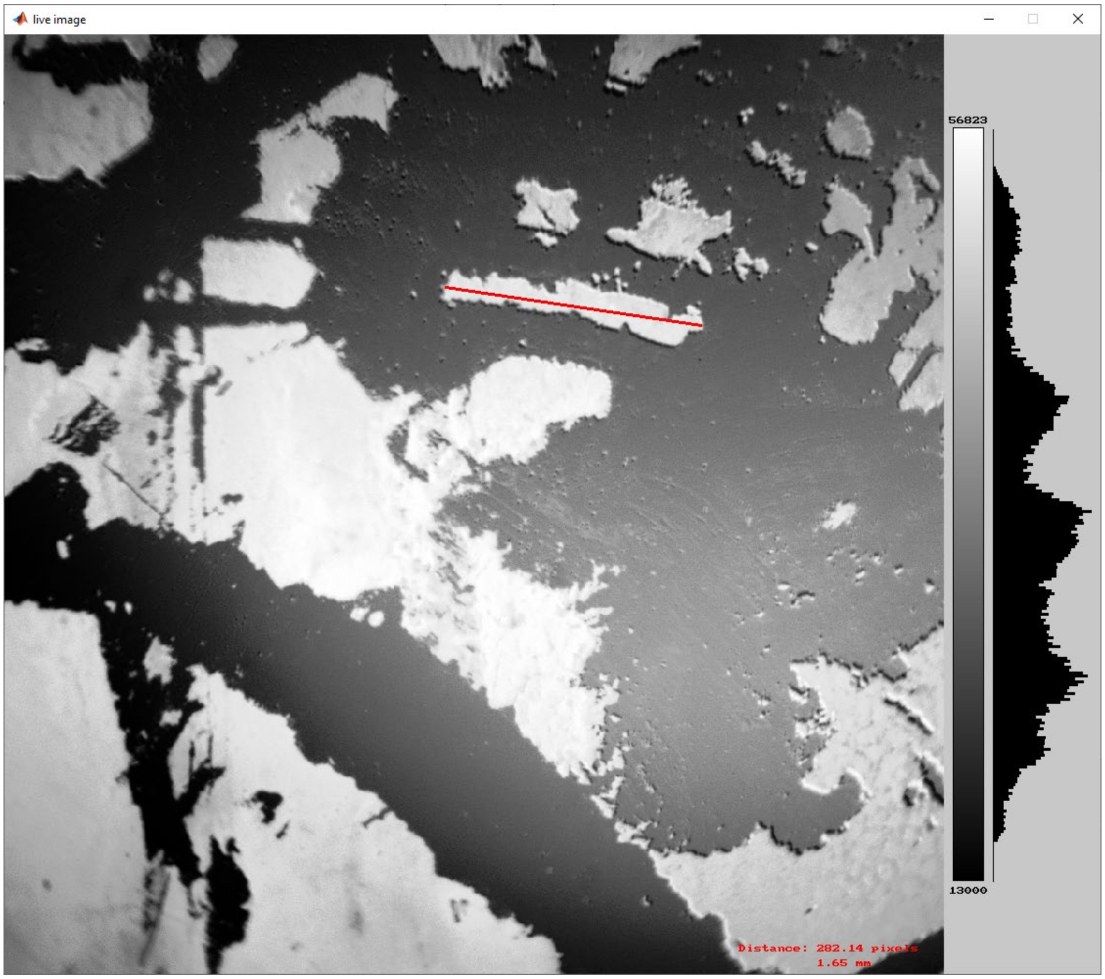


**Supplementary Figure 3: Real-time camera display.** Separate camera display with color bar and histogram. Ranges can be toggled between fixed (set by user) or automatic (min to max pixel value). The displayed histogram can be used to maximize contrast by applying histogram equalization. Different color maps are available (grey, grey inverted, hot, jet). The stream can also be used to measure distances between points and to select ROIs for zooming or displaying in the main tab.

a

b

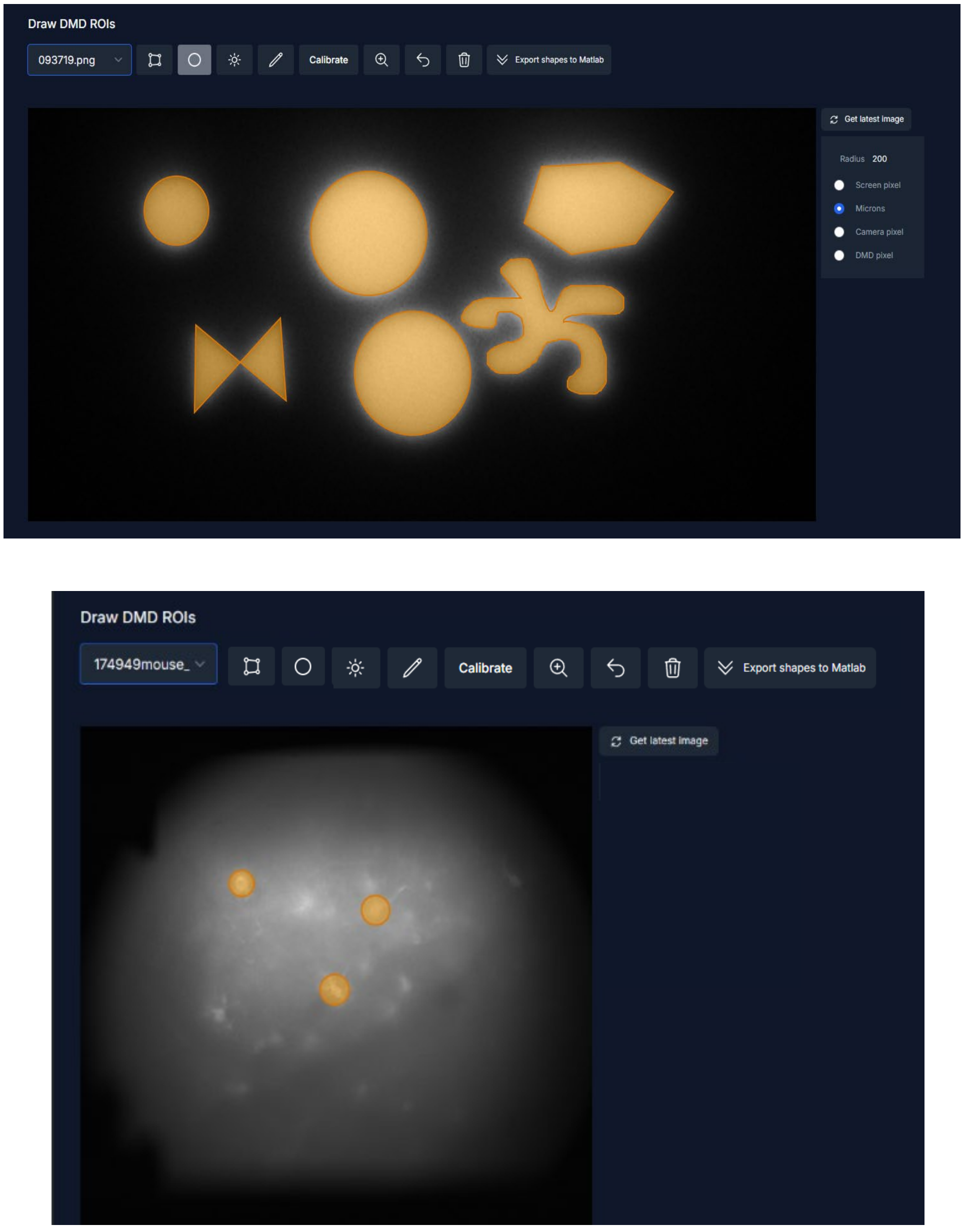


**Supplementary Figure 4: Luminos patterning tab.** The patterning tab (a DMD tab in this case) provides tools for drawing target regions of interest. The user can then load a reference image using the dropdown selector, and can draw arbitrary polygons, circles, or curves. The pattern is uploaded to the DMD automatically. The DMD tab does not currently provide the ability to load multiple patterns into a pattern stack in DMD memory, but this capability is present in the MATLAB code and may be easily scripted into an experimental protocol. a) Demonstration of DMD drawing modes: Polygons, freehand curve and circles. Circle sizes can be set in multiple units, including µm (by setting magnification in objectives panel in main tab), camera pixel or DMD pixel. b) DMD pattern drawing in widefield snap of mouse brain, here three circles for selection of individual cells.

**a**

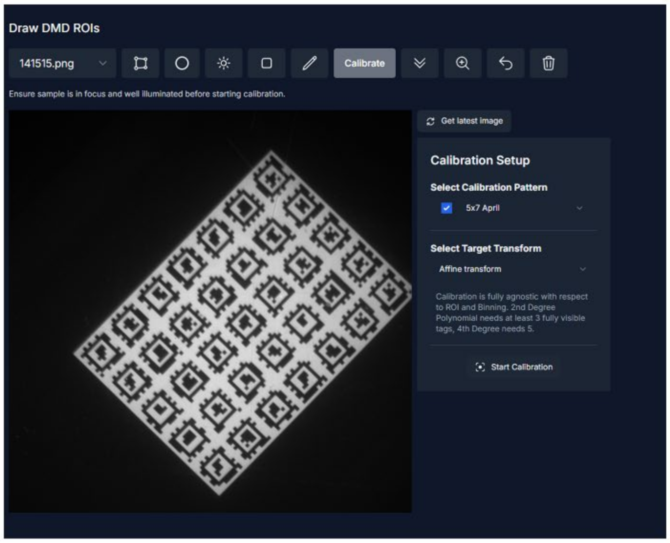


**b**

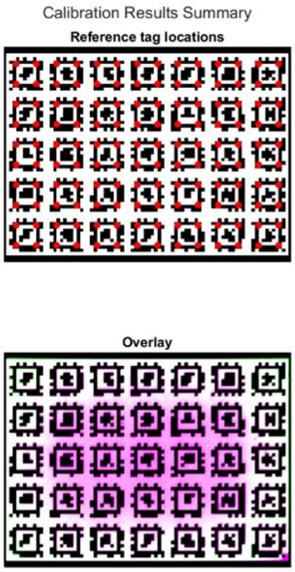


**c**

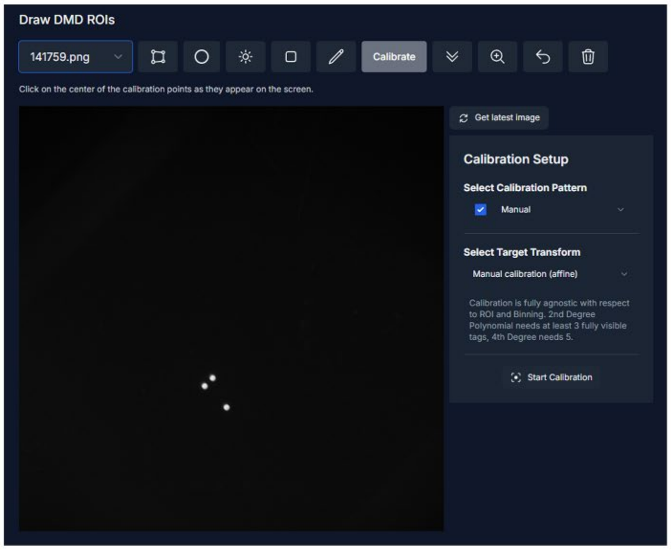


**Supplementary Figure 5: Luminos patterning device calibration.** a) DMD calibration tab. Selection and preview of arrays of April tags for automated DMD calibration and target transformation between camera and DMD coordinate systems. April tag pattern can be selected from the dropdown menu. Preview of the pattern can be toggled to focus on the sample (typically fluorescent slide or glass slide with fluorescent liquid) and find a homogeneous area. Upon pressing the “Start Calibration” button, Luminos automatically detects tag corner positions, and updates/saves transform. Different tag sizes are available to use with different microscope illumination areas. Manual calibration option (by clicking on three sequentially displayed points) is also available. b) Results of a calibration. Top: DMD pattern with detected April Tag corner points marked in red. Bottom: Overlay of projected DMD pattern and transformed camera image, showing excellent registration. c) Pointwise calibration using three sequentially projected points (number and position can be set in Rig initializer file). Automatic pointwise calibration is used for galvos and SLMs, and is available as a manual option for DMDs (clicking on points as they appear).

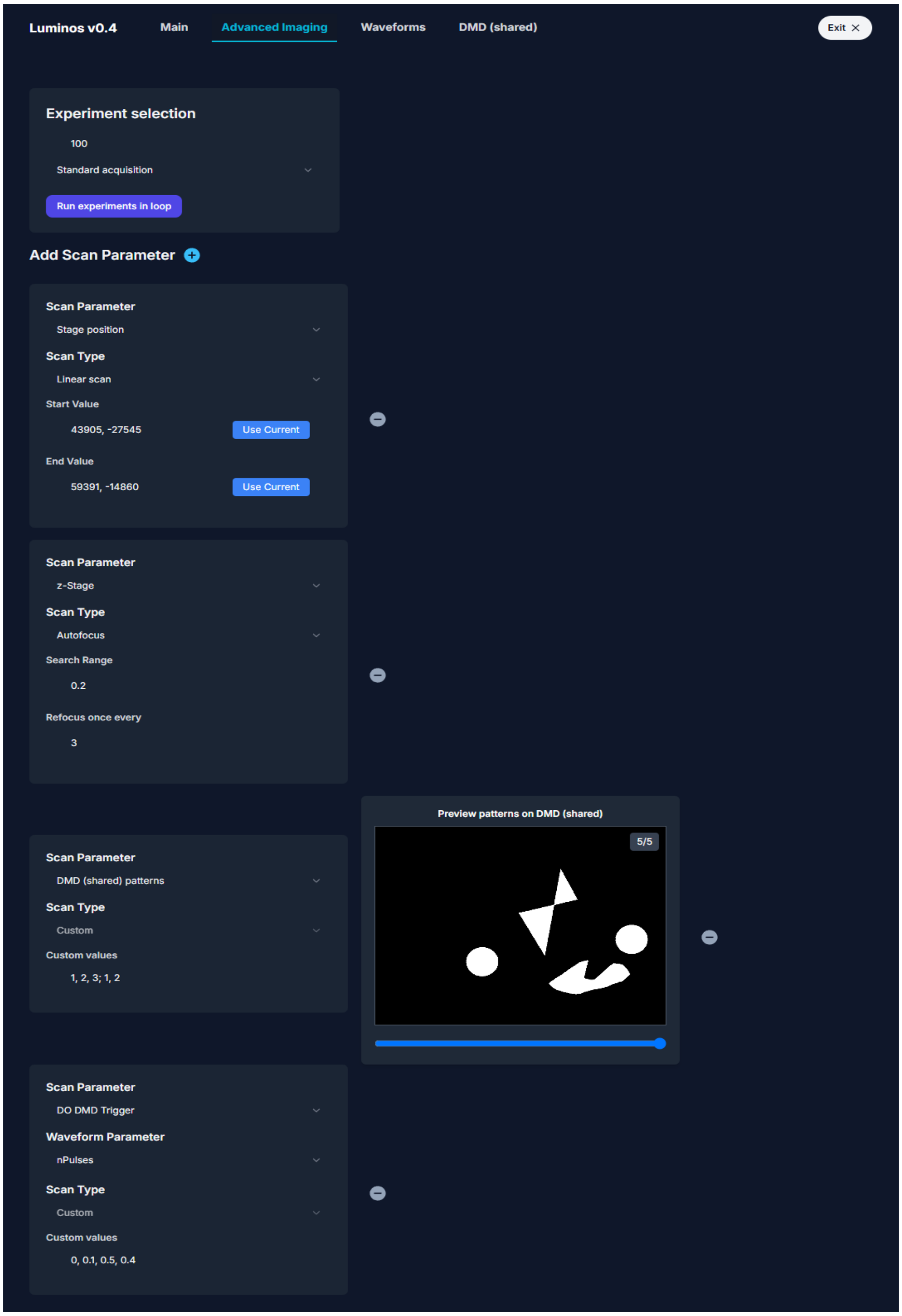


**Supplementary Figure 6: Multi-round tab.** The Multi-round tab can be used to set up multi-round experiments. Experiment selection: Pick number of repetitions and type of experiment. Options for Hadamard (with selection of Hadamard pattern parameters) and HiLo structural imaging appear if DMDs are available. Use Scan Parameter: Vary parameters of connected stages (e.g. for z-Stacks, automated multi-well plate acquisition), parameters of generated waveforms from waveform tab (e.g. varying stimulation intensity, pulse width), or DMD patterns between experiments, in arbitrary combinations.

```
Command Window
>> app

app =

  Adaptive_Upright_App with properties:

                     Devices: [1×21 Device]
    monitored_devices_index: []
                explistener: [1×1 event.listener]
                 Experiment: 'Optopatch'
                  expfolder: []
                   datafile: []
                   basepath: 'C:\Updated_Control_Software\luminos-private\'
                 datafolder: "D:\Phil Brooks\Optopatch\20240405"
              server_target: "X:\Lab\Labmembers\Phil_Brooks\Data\Optopatch\20240405"
                   Rig_Init: [1×1 Rig_Initializer]
                       User: [1×1 User_Key]
                    logfile: "D:\logfile.txt"
             screen_blanked: 0
               exp_complete: 0
             isDataAcquired: 0
                   jsServer: [1×1 JS_Server]
                      VR_On: 0
                   VRclient: []
        wasAppDeletedFromJS: 0
                     jsPort: 3010
                    gitInfo: []
                    rigName: 'Adaptive_Upright'
                       tabs: [1×6 string]

>> cam = app.getDevice("Camera");
>> cam.exposuretime

ans =

    0.1000

>> cam.exposuretime = 0.05;
>> cam.exposuretime

ans =

    0.0500

fx
```

**Supplementary Figure 7: MATLAB command window interaction with Luminos.** All aspects of the Luminos app can be controlled via MATLAB command line. The Luminos app exists as an object in the MATLAB workspace and can be interrogated and controlled directly from MATLAB either through scripting or through live commands. Here, we display a summary of the app state. We extract the camera device, query the current exposure time, change the exposure time, and then check that this change was effective. The updated exposure time will be reflected in the UI camera panel as well.
